# Torsion in motion: the visual system as a three-axis gimbal

**DOI:** 10.64898/2026.08.09.743586

**Authors:** A.H. Méndez, J. Otero-Millán, C. de la Malla, J. López-Moliner

## Abstract

Eye movements are usually characterized by where we look and represented using the two rotational degrees of freedom that direct the line of sight onto a target. The third, torsion about the line of sight, is often regarded as a limited response and has rarely been measured outside the laboratory. Yet the conditions producing the largest torsional responses, looking at a target straight ahead, are the ones that dominate natural walking. Here we tested whether torsion is present during this pervasive behavior and measured its impact on the visual input. Ten participants walked toward a distant target wearing a mobile eye tracker fitted with an iris camera to estimate torsion. Comparing iris feature rotations in the eye camera with simulated projections of candidate three-dimensional eye rotations, eye rotations were poorly explained without torsion and best explained by substantial torsional counter-rotation to head roll (gains .6–1). In the reconstructed retinal flow, this torsion cancelled the rotation added by head roll, leaving flow dominated by the observer’s translation through the scene. Together, these results reveal that torsion is a substantial component of eye movements during locomotion and show that realistic models of oculomotor control and visual input during self-motion must account for all three of the eye’s rotational degrees of freedom.

## INTRODUCTION

Eye movements are typically framed as changes in where we look, yet where we look does not tell us how the eye rotates in three dimensions. Geometrically speaking, infinitely many rotations can carry the line of sight between two positions. Figure 1A illustrates two pupil positions and two rotations that could shift the pupil between them. Any rotation described by an axis that lies on the plane shown in Figure 1A can do the same job. Gaze direction alone thus leaves one rotational degree of freedom, torsion, unconstrained. Yet, in many cases, neglecting true torsional components can be a well-motivated simplification. Because torsion is the component of eye rotation about the line of sight, it does not generate displacements at the fovea (see Figure 1B). Gaze stability can therefore be described without torsion (Grossman et al, 1989; Crane & Demer,1997; Dietrich & Wuehr, 2019). In other cases, omitting torsion is a good approximation of the eye’s 3D rotation. This is true when the eye makes a saccade or smooth pursuit in head-restrained conditions near its forward resting position, a common setting in controlled studies, and in which the first laws of eye movements were described (von Helmholtz, 1867). According to Listing’s law, all observed eye orientations can be described with respect to a reference orientation (the primary position) by rotations whose axes have no components about the line of sight (van den Berg, 1995). This constraint on orientation has implications for how the eye actually rotates in space. Between any two orientations, the rotational axis - and its torsional component – are constrained to a plane (displacement or half-Listing’s plane) that tilts with eye eccentricity (Tweed & Vilis, 1990, see Figure S1). As a consequence, saccades and smooth pursuits from and towards the primary position have no torsion. Those occurring between targets near the primary position have minimal torsional components (see section Three-dimensional eye rotations, torsion and Listing’s law on SI for a more detailed description). Taken together, both an interest in gaze stability and the study of eye rotations in head-restrained conditions can be reasonable reasons not to account for torsion in the study of eye movements, but neither implies that everyday eye rotations truly lack a torsional component.

**Fig 1.**
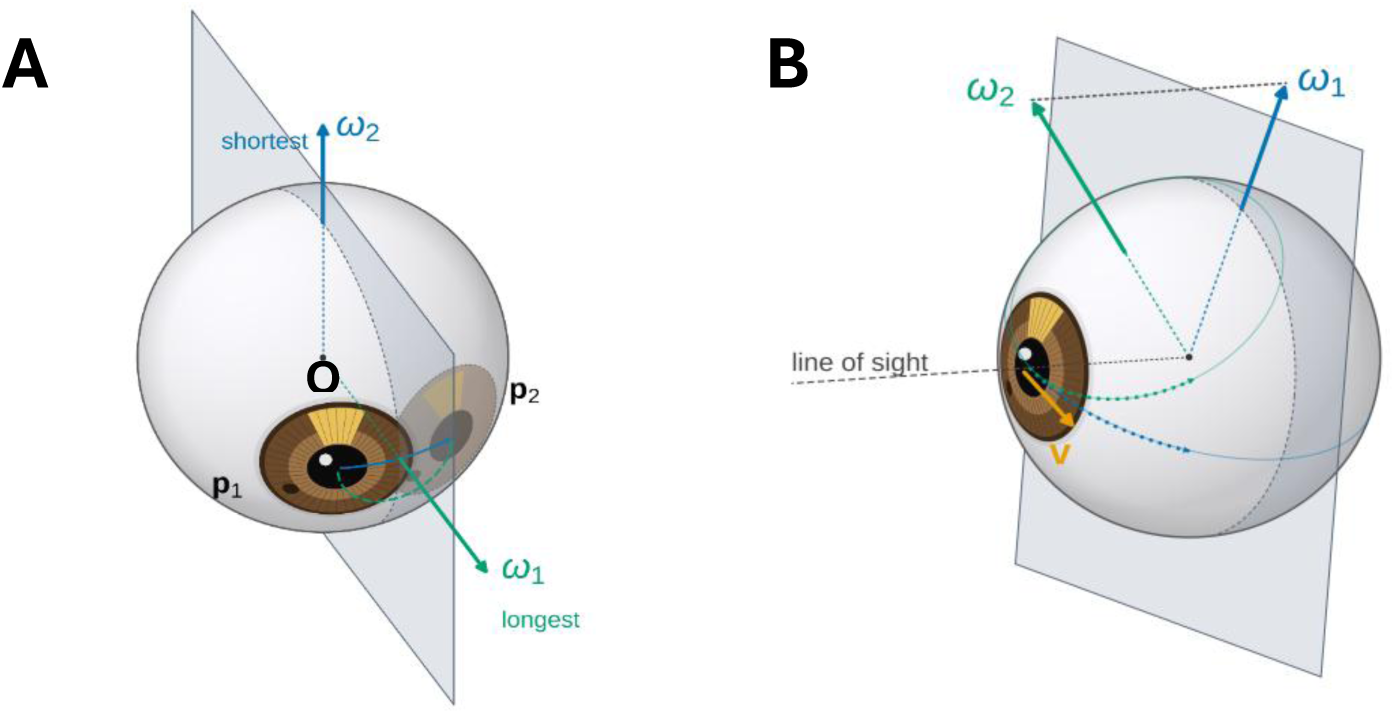
*A.* Two different rotational displacements **ω (or velocities)** that rotate the pupil **from P_1_ to P_2_**, using the right-hand rule. Notably, the rotational displacements chosen are the ones with the longest (**ω_1_**, green) and shortest arc (**ω_2_**, blue). There are infinite rotations that rotate the pupil between those two positions, and they all live in the plane that bisect **OP_1_**and **OP_2_**, being O the eye center. Of all these rotations, only **ω_2_** (blue) explains the rotation of the pupil and all iris features. That is, only one fixed axis orientation explains the rotation between two different eye orientations. *B.* Rotations that differ in their component along the line of sight generate the same instantaneous displacement on the fovea (infinitesimal step). **ω_1_** (blue) lies on the plane perpendicular to the line of sight, so carries no torsional component. ω_2_ (green) differs from ω_1_ only in its torsional component. The linear velocity shown is tangent to the sphere, orthogonal to both **ω** vectors and parallel to the plane. The orange vector is not an angular velocity, but the instantaneous linear displacement of the fovea caused by the eye rotations.

The need for three degrees of freedom to fully describe eye kinematics outside the laboratory depends on how prevalent torsion is during natural behavior. Controlled studies of eye movements exposed to head rotations show that – although limited in range - the eyes can clearly counter-rotate around the line of sight to stabilize the visual image as a consequence of the vestibulo-ocular (VOR) reflex (Raphan & Cohen, 2002). When a target is visible, the mechanisms underlying the control of eye torsion combine VOR and optokinetic (OKR) responses. During the OKR, rotational patterns produce rotations of the eye about the line of sight in the same direction as the pattern, with gains not larger than 0.2 (Cheung & Howard, 1991; Farooq et al, 2004). VOR and OKR work synergistically to stabilize visual input (Wibble et al, 2022). Scleral coil recordings show that the largest torsional gains, in response to a 10-degree active head roll, occur in the presence of a visual target and at 0.6 and 1.33 Hz, with average gains of approximately 0.6 and 0.7, respectively (Collewijn et al, 1985). Two studies analyzing the interaction between eye orientation (at 15 degrees from primary position) and the VOR, reveal low gains in eccentric positions and bigger values of up to 0.5 when the eye is in primary position in the dark (Tweed et al, 1994a) and 0.6 while stabilizing a visual target (Fetter et al, 1995). According to the authors, these results suggest that the torsional VOR primarily serves foveal rather than whole-retinal stabilization, and reflect a compromise between image stabilization and compliance with Listing’s law (Misslich et al, 1994, Tweed et al, 1994b). Importantly for our work, the largest torsional gains described occur when the head rolls in conditions frequently met during locomotion: a visible target with the eyes near the primary position. Studies using head-mounted eye tracking technology show people tend to explore the world with head and eyes aligned (Hollands et al, 2002; Hart & Einhäuser 2012), with standard deviations around 10 degrees across both the horizontal and vertical directions (Foulsham et al, 2011). By contrast, how humans rotate their heads around the roll axis while walking has received considerably less attention. Prior studies of head and eye stabilization during locomotion have predominantly characterized only their horizontal and vertical components (Grossman et al, 1989; Pozzo et al, 1990; Moore et al, 2001; Dietrich & Wuehr, 2019). Consequently, if participants do roll their heads during locomotion and the eye behaves as in controlled studies, torsion could compensate for approximately 60-70% of the head’s roll component and an accurate account of eye rotation during locomotion would require all three of the eye’s rotational degrees of freedom.

How the eyes rotate in 3D to stabilize the retinal input throughout the gait cycle, when proprioceptive, visual and motor cues are simultaneously available, remains an open question. There is still no widely accepted non-invasive method to accurately measure the full 3D rotational kinematics of the eyes in the head outside the lab (see Otero-Millán et al, 2015 for a review, but see Imai et al, 2001). Recovering ocular torsion from video alone is confounded: the apparent rotation of iris features on a head-fixed camera reflects not only torsion but also a geometric projection artifact that scales with gaze eccentricity and its relation to the axis of rotation. However, stabilizing gaze at the front is not only a highly frequent behavior but represents a convenient framework in which to estimate torsion (see Eye stabilization in natural behavior on SI). Research shows that at viewing distances exceeding one meter, eye stabilization is achieved through counter-rotations of the eyes relative to the head (Moore et al. 1999; Moore et al., 2001) with a precision of about 0.1 degrees (Grossman et al, 1989), allowing for accurate estimations of gaze in the head (see 3D gaze vectors and gaze stabilization on SI). Knowing gaze substantially constrains the candidate eye rotations that can explain eye measurements from a camera image. Torsion can then be estimated from computing the 3D candidate eye rotation that best predicts the iris feature rotations measured on the image, and (see Torsion estimation on Methods).

Characterizing the full 3D eye rotational kinematics is not only fundamental for the study of oculomotor control but also for building accurate models of the stimulus that reaches the retina. Advances in head-mounted recording now make it possible to measure this stimulus in naturalistic settings, in freely moving rodents (Wallace et al., 2025) and in zebrafish in their natural habitats (Alexander et al., 2022). In humans, wearable sensors record gaze – not 3D eye kinematics-, head and body movements simultaneously (Kothari et al., 2020; Ma et al., 2024; Greene et al., 2024). Because torsion rotates the image about the fovea, small torsional angles will have small effects on where stimuli fall on the retina. The rotational velocities needed to counter head roll at every step, however, need not be small, and their effect on image motion grows toward the periphery. How torsion shapes the retinal flow available during locomotion is a matter that informs a long-standing debate over how the brain uses flow cues to recover and control self-motion (Lappe & Van den Berg, 1999; Warren et al 2001). Current egocentric studies of retinal flow during locomotion suggest fixations represent an ideal window for the extraction of self-motion parameters (Durant & Zanker, 2020; Matthis et al, 2022; Zorpala & López-Moliner, 2026). These studies – as the human databases cited above – report gaze direction as a proxy for eye 3D rotation. Thus, the contribution of torsion to the retinal flow available during locomotion remains unexplored.

Here, we leveraged head-mounted eye tracking technology (Figure 2A top), computer vision algorithms and a simulation-based framework linking eye kinematics to their camera projection to test whether torsion is present during walking and how it shapes retinal flow. We measured eye rotations as participants walked towards a distant target (from 14 to 4 meters away from the target). For each pair of frames we: i) used the gaze and head data to simulate a set of physiologically motivated candidate 3D eye rotations – a torsion-free scenario, a scenario with rotations predicted by Listing’s law, and multiple scenarios with head-counter-roll (VOR) rotations each with a different torsional gain (0 to 1) –, ii) projected each through the eye-camera geometry, and iii) identified which scenario produced feature rotations that best matched the ones measured in the real camera image - tracked using CoTracker, a deep learning-based point-tracking model (Karaev et al, 2024, see supplementary videos 1 and 2). In other words, torsion was estimated as the component of rotation about the line of sight that best explained the feature rotations observed in our device’s eye camera. We then used the measured head and eye kinematics to estimate the influence of torsion on retinal flow. At any point on the retina, the flow generated from self-motion can be resolved into its translation and rotational components (Longuet-Higgins & Prazdny, 1980). In our study, as participants fixated on a distant target, forward translational flow would resemble expansive patterns and non-compensated head roll components would result in flow curl (see Tweed et al,1994b), simplifying the interpretation of torsion’s role in shaping retinal flow. Finally, because non-compensated head roll can interact with the curl generated by the projection of the surface where gaze is stabilized, a cue proposed to guide stepping on complex terrain (Matthis et al., 2022) and heading (Zorpala & López-Moliner, 2026), we modeled both sources of curl together to study their interaction.

**Figure 2.**
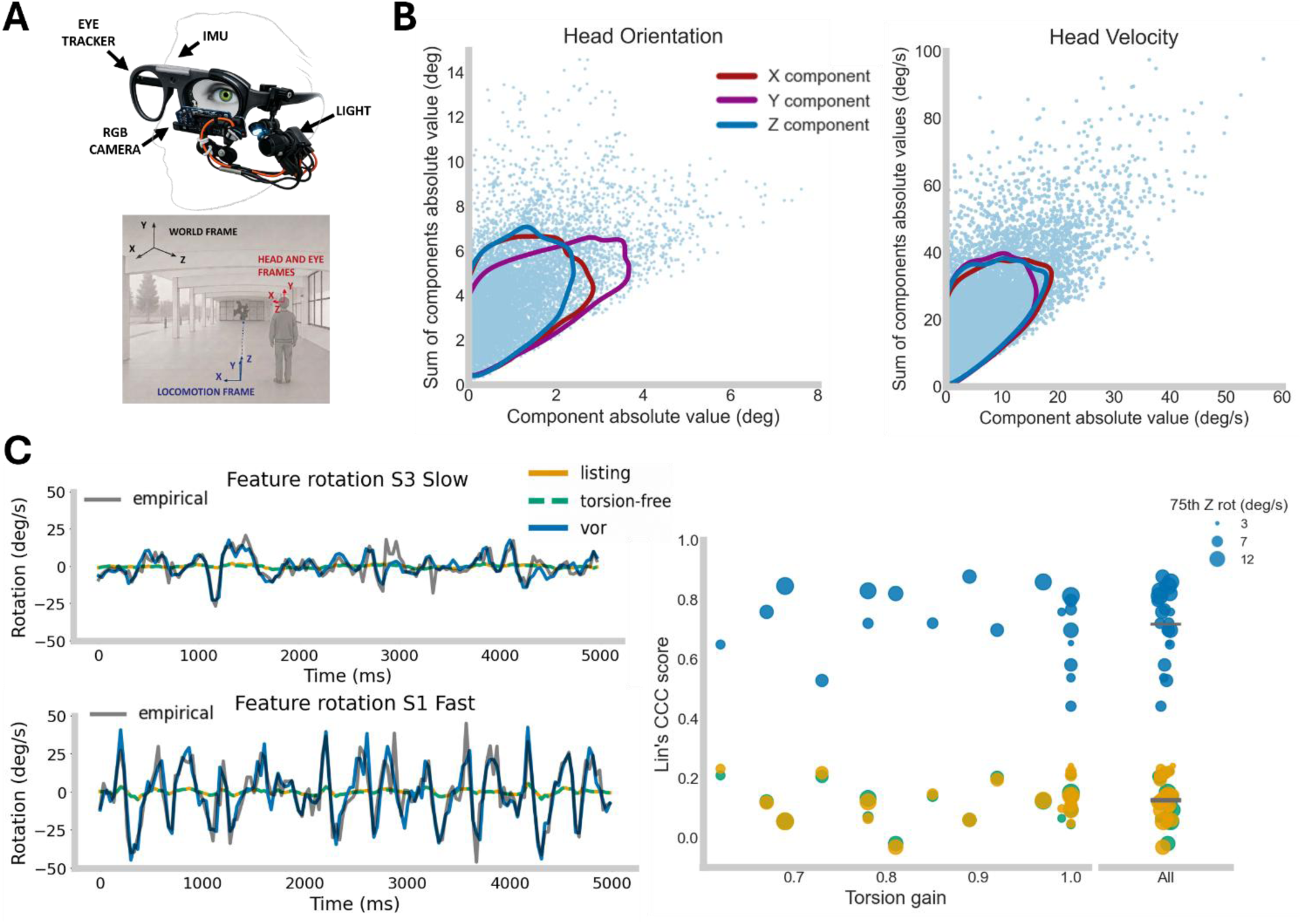
***A.** Top.* Custom-made head-mounted device combining the Neon eye tracker (Pupil Labs), an RGB camera and a dimmable light. *Bottom.* Four 3D frames of reference (FoR) are relevant for this study, two static (world and locomotion) and two subject centered (head and eye). The Z axis of the locomotion, head and eye FoRs are approximately aligned throughout the trial. For the locomotion FoR the Z axis is fixed in the world and points forward (towards the target). The head’s Z axis moves with the head but – as subjects are fixating a target along their path -, it also points approximately forward. The eye’s Z axis also moves with the head and its exact forward orientation will depend on compensatory eye movements. ***B.*** *Left.* Blue dots represent the Z component of the head’s orientation vector (on the locomotion frame) on the X axis, and the sum of all three components on the Y axis; for each frame for all corpus data. Blue contour is the 75th percentile 2D density distribution of the blue dots. Red and violet contours represent the 75th percentile for the X and Y components of head orientation, respectively. *Right*. Same logic but applied to the head’s velocity vector. **C.** *Left.* Two examples showing the mean rotation of iris features over the course of a slow (top) and fast (bottom) trial. Colored lines show each of the 561 simulated cameras for a given scenario (one color per scenario); the black line shows the camera from the empirical data. *Right.* Trial-level mean fit score of each scenario with the empirical data is represented as a function of each subject’s fitted gain. 20 dots represent 10 subjects x 2 trials. On the rightmost column, all values are aligned vertically to show the mean fit score across trials for the three scenarios. Size indicates the 75th percentile of head z component for each trial.

## RESULTS

To first describe head 3D rotational kinematics, we analyzed the inertial measurement unit (IMU) recorded from subjects’ forehead (Figure 2A top) as they walked along a straight path on two occasions - at a slow and a fast pace - while fixating a target (Figure 2B bottom; also see supplementary videos 1 and 2). Viewing distance decreased from 14 to 4 m over the course of each trial. The two paces were introduced to increase gait variability. For this reason, results from the fast and slow trials were combined. Separate analyses of each pace yielded qualitatively similar results, and pooling therefore did not affect the conclusions of the study. Figure 2B shows the magnitude of each component of the head’s orientation and rotational velocity vectors, pooled across the entire corpus (all frames, subjects and trials). Each component is compared against the sum of all three components for each frame to appreciate its relative contribution. The head orientation data reveals that the head displacement from baseline is small and rarely exceeds five degrees in any component (75th percentile: x = 1.82 deg, y = 2.49, z = 1.63). Velocity data shows that the head rotational dynamics imposes similar demands across all three axes of rotations. Subject-level analysis shows inter-subject differences in the components of both the head’s orientation and velocity vectors (see Figure S2A). These results indicate that head roll during locomotion is non-negligible, such that compensatory eye counter-rotation must include a torsional component.

To measure torsion, we simulated the candidate eye rotations that could explain how subject’s stabilized gaze on the target and selected the torsional value that best explained the feature rotations observed in our device’s eye camera. For each subject, camera position and orientation were adjusted to secure high-quality iris imaging. To estimate how much uncertainty in our measured camera position relative to the eye affected our torsional estimates, we repeated simulations across 560 camera positions, varying position by ±1 cm in all directions and roll by ±10° (see section Camera Uncertainty on SI). In addition, 3D gaze data for the simulations was derived assuming the eyes fully compensated for horizontal and vertical head displacement to keep gaze on the target - an assumption both supported by prior reports (Grossman et al. 1989) and by our gaze and image data, which showed that participants stabilized gaze with an average root mean square (RMS) dispersion of 1.62° (between-subject SD = 0.67°) across all trial data and 0.74° (between-subject SD = 0.44°) within the 5-second window of each trial with strongest stabilization that was selected for our analysis (see section Gaze 3D vectors and gaze stabilization on SI).

Figure 2C summarizes the results for three of the simulated scenarios: no torsion, torsion as described by Listing’s law and the VOR torsional gain that best fit the data. Figure 2C left shows the temporal relation between the empirical rotation of features and those predicted by each scenario, for two trials and all simulated cameras (plots for all time-series are available, see Data availability). Concordance analyses showed the VOR scenario best explained the data (Figure 2C right), with an average gain of 0.88 across subjects (see Figures 2C and S2C; gain std= 0.13; mean and std for CC scores: torsion-free = 0.12 +/- 0.07, listing = 0.13 +/- 0.07; VOR = 0.72 +/- 0.13). Across subjects, the lag that maximized the correlation between the feature rotations from our camera and the best fitted VOR scenario across all participants was −1.66 ms. (std = 17 ms.; mean correlation = 0.75, std = 0.10). As expected, the small range of head oscillations while subjects stabilized gaze on a distant target rendered our torsion-free and listing scenarios almost identical. On average, the gaze vector was 2.4 degrees from its baseline, yielding a difference of only 1.2 degrees between the torsion-free and half-Listing’s-law planes at each gaze position. In other words, our empirical data is explained by compensatory eye rotations with significant torsional components. Together with the head data, this reveals the pervasive presence of eye torsion as participants walked towards a target and confirms the need for a 3D description of eye rotations to accurately model eye behavior during locomotion.

To estimate the relevance of measuring torsion for models of retinal flow, we then calculated the head-centered and retina-centered flow generated by the torsion-free and fitted VOR scenarios, assuming perfect gaze stabilization (see Retinal flow and curl estimation section on Methods). For each pair of consecutive frames, we computed both the optic flow and curl magnitude (maximum curl value across the vector field). The relation between curl, gain and head roll rotation can be clearly seen in Figure 3A, both at corpus (left) and subject level (right). Figure 3B describes the retinal flow of specific frames with varying head roll strength and gain. Altogether, results show torsion compensates for the head-generated flow curl during locomotion. In this data, torsion’s effect on flow is negligible only in frames with no head roll velocity (Figure 3A left and 3B top row) and subjects that do not roll their heads while walking (Figure 3A right, see head variability across subjects in Figure S2A).

**Figure 3.**
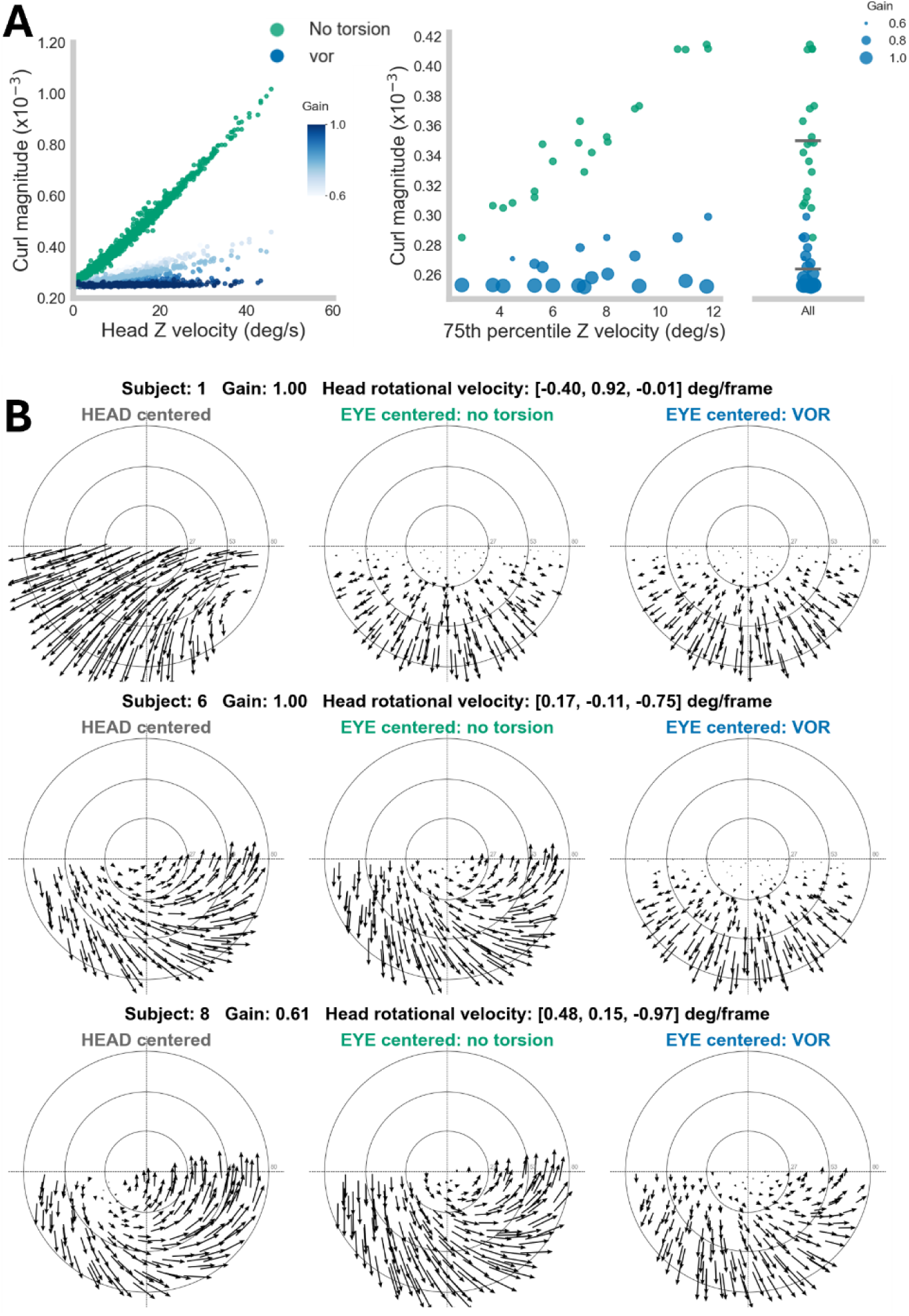
**A.** *Left.* Each dot represents the max curl magnitude of the simulated torsion-free (green) and best fitted VOR (blue) retinal flow for each frame of the corpus data. Blue VOR is shaded according to the subject’s gain. *Right.* Each dot represents the trial level mean curl for the torsion-free (green) and VOR (blue) scenarios for a trial, as a function of the 75th percentile head’s angular velocity Z component. 20 dots represent 10 subjects x 2 trials. On the rightmost column, all dots are aligned to show the mean for each scenario across trials. **B.** Rows represent the head-centered (left) and eye-centered torsion-free (middle) and VOR (right) flow for one frame (details in each title). A fixed linear speed of 1 m/s to allow direct comparison across subjects and trials. The top row represents an instant where the head has little Z rotational velocity component, with almost no curl in both stabilized torsion-free and VOR scenarios. In the middle and bottom rows, head rotational displacement has a predominant Z velocity component, and therefore the point of minimal velocity in the flow field is almost at the center for all three panels. When gain is high (middle panel) eye torsion clearly decreases rotational curl. Intermediate gains (0.64, low panel) decrease curl, but curl is still visible to some extent in the VOR scenario.

Finally, we simulated the role of torsion in mediating the interaction between roll-generated curl and curl generated by the relative motion of the observer with respect to a laterally fixated target (as in Matthis et al 2022, and Muller et al, 2023). The sign of this latter source of curl depends on the side of the fixation point relative to the subject’s trajectory and could inform the control of stepping (Matthis et al 2022). To study this interaction, we extracted the 50^th^ and 75th percentile values of head roll and the median torsional gain from our data. Figure 4 top shows that these non-compensated values of the head’s rotational velocity about its roll axis (Z) expected during locomotion can both stabilize and reverse the curl generated by the fixation plane on the retina. In other words, for flow cues generated from the geometry of the fixation plane to emerge, the eye needs to be stabilized across all three degrees of freedom (Figure 4 bottom). A complete model of the incoming retinal flow during natural behavior will require accurately tracking each subject’s unique pattern of the head’s translational and rotational kinematics (see Eye stabilization in natural behavior on SI).

**Figure 4.**
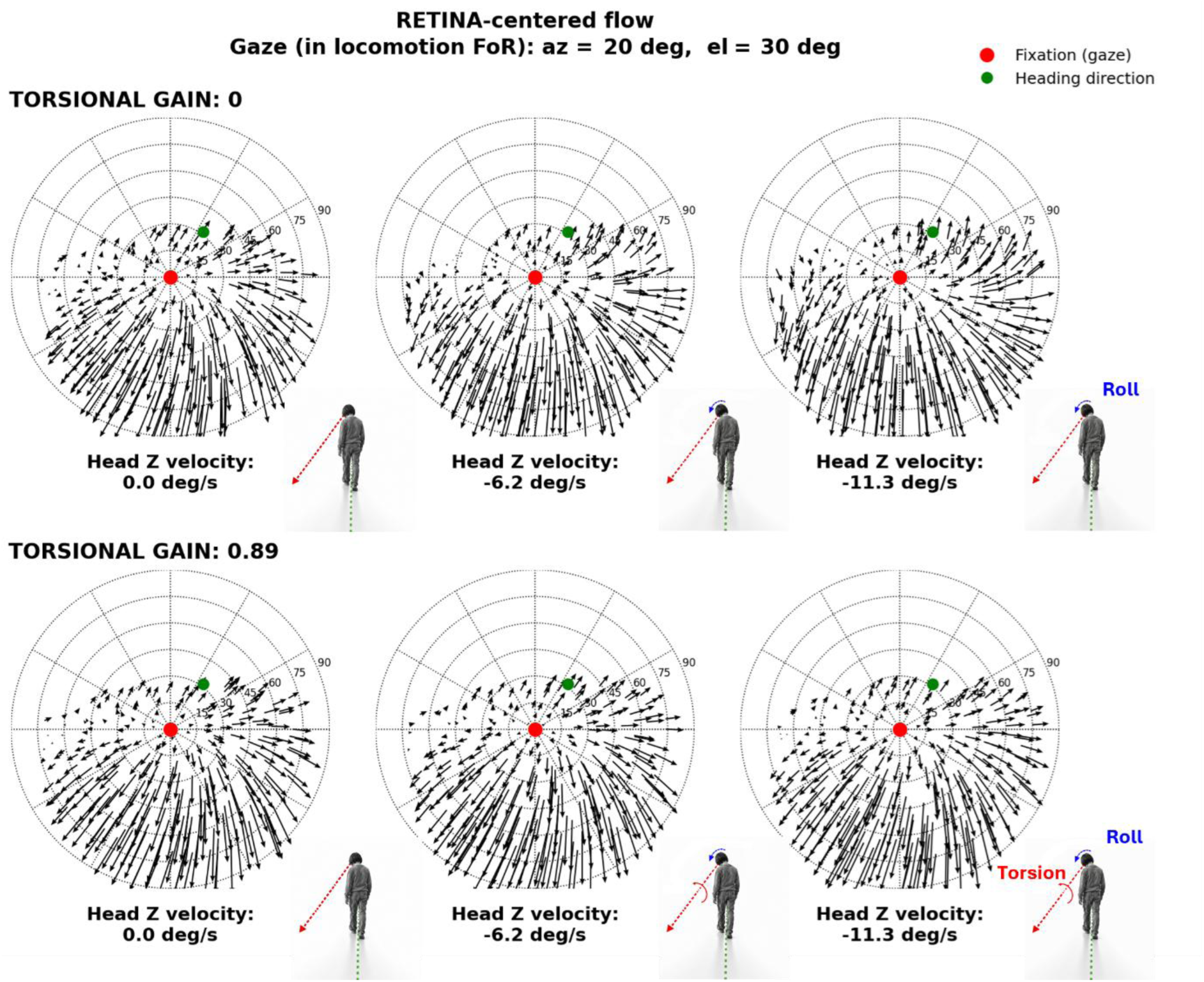
Retinal flow generated while walking and stabilizing gaze on the floor and laterally. For all six panels, the person starts looking with head and eyes aligned on the target at the floor and, on the next frame, walks forward while rotating the eye to stabilize the flow at the fovea. Each column varies in the head’s rotational velocity about the Z axis (roll component). Each row varies on the eye’s torsional gain. Top row corresponds to a VOR gain of 0 – stabilizing the fovea without torsion - and bottom row to the average gain obtained from our data (0.88). When the head has no rotational velocity about its Z axis (left panel, same for both rows), flow curl emerges from the relative motion of the fixation plane relative to the moving subject (as in Matthis et al, 2022). Middle and right panels show two different head-centered rotational velocities about the Z axis, extracted from our walking data (50th and 75th corpus percentile values). The top row shows how head roll during walking stabilizes (middle panel) or changes the sign (right panel) of the flow curl seen in the left panels. Bottom row middle and right panels show how torsional eye movements with high gain can recover the curl available in the left panels. Walking speed is 1 m/s and forward. The gaze position in the retinal flow is at the center (red dot). The walking direction is represented by the green dot (and the green trajectory in the inset figure).

## DISCUSSION

The current study addressed a dimension of head and eye kinematics that has been largely overlooked — rotation around the forward axis (roll) and the line of sight — and showed that torsion is not a limited response but a substantial component of eye movements during locomotion, with systematic effects on retinal flow under naturalistic conditions. The study of torsion has been both a methodological and conceptual challenge (see Three-dimensional eye rotations, torsion and Listing’s law). Here, we leveraged head-mounted technology to capture high-quality iris images, which together with gaze data and kinematic simulations allowed us to test competing three-dimensional eye rotations outside the lab.

Roll and torsion are intrinsic components of head and eye kinematics when fixating a target during locomotion. Head roll during walking, largely unexamined in previous studies of locomotion (Grossman et al., 1989; Pozzo et al., 1990; Moore et al., 2001), was small in amplitude but comparable in velocity to rotation about the other two axes, so that compensatory eye movements necessarily required a torsional component. This is confirmed by the pervasive presence of torsion in our data, highlighting the need to account for torsion to rigorously describe eye kinematics in natural behavior. The high gains reported here – .88 on average and ranging from .61 to 1 – are consistent with the larger torsional gain values reported by previous studies (Tweed et al, 1994a; Fetter et al, 1995). Our large values could be explained by the expected inter-subject variability (Figure S2C), the activation of the torsional VOR with the eyes near primary position – where torsional gain is largest - , and by differences in head dynamics during locomotion from the larger, fixed-axis head rotations used to characterize torsion in controlled studies (e.g., Collewijn et al., 1985). In addition, feed-back and feed-forward signals available during active locomotion may further increase torsional gain (Ibbotson et al., 2005; Lambert et al., 2023; Wibble et al., 2024). Considering that people often explore the world with eyes and head aligned (Hollands et al, 2002; Hart & Einhäuser 2012), our results suggest that conceiving torsion as a limited response is an underestimation of its true role in natural behavior in humans. As our ability to measure torsion non-invasively and in larger samples increases, a clearer picture of inter-subject variability in torsional responses will emerge.

Our data reveals the need to account for torsion when analyzing the structure of retinal flow generated by self-motion. Retinal flow combines a translational component, which carries information about heading and scene layout, and a rotational component, which does not (Longuet-Higgins & Prazdny, 1980). By compensating for head roll, torsion removes rotation about the line of sight, leaving flow dominated by the observer’s translation through the scene. In this sense, the eye behaves as a three-axis gimbal, stabilizing not only the direction of gaze but, to a large extent, the orientation of the retinal image relative to the world. Discussions on the use of retinal flow in navigation have revolved around the computations that allow relevant information to be extracted from flow fields (Llewellyn, 1971; Warren & Hannon, 1990; Burlingham et al., 2020), while others have centered on the complexity of the incoming flow during natural behavior (Calow & Lappe, 2007). Current egocentric studies suggest that retinal flow is used during periods of gaze stabilization (Durant & Zanker, 2020; Matthis et al., 2022). These studies, however, do not include torsion when reconstructing retinal flow. Our results suggest that the flow reaching the retina during locomotion is more stable than retinal flow reconstructed from head-camera images and two-dimensional gaze. How these more stable patterns of flow contribute to the extraction of heading during natural locomotion remains to be tested empirically (but see Zorpala & López-Moliner, 2026). Our results show, however, that cues generated by fixation, such as the curl proposed to guide foot placement (Matthis et al., 2022) and heading (Zorpala & López-Moliner, 2026), can be masked or even reversed by head roll unless torsion compensates for it (Figure 4).

Whether studies neglecting torsion have misrepresented retinal flow depends on the presence of head roll and the frame of reference in which eye rotation has been specified. In our data, torsion and its consequences on flow were negligible only in frames without head roll velocity, or in subjects who do not roll their heads. Across very different approaches - including analytical models of natural flow statistics, simulations of retinal flow during locomotion, flow-based models of heading and object-motion detection, and synthetic stimuli for training neural networks - , eye (or camera) rotation is often specified relative to the world, with torsion omitted or eye rotations derived from the requirement to stabilize the fovea (e.g., Calow & Lappe, 2007; Mineault et al., 2021; Maus & Layton, 2022; Lutwak et al., 2025; Zorpala & López-Moliner, 2026; see also the simulations in Matthis et al., 2022). Because foveal stabilization leaves rotation about the line of sight unconstrained, a natural and parsimonious choice is to set it to zero. Our results suggest neglecting torsion in the world approximates the retinal input well, since no torsional rotation of the eye relative to the world produces flow similar to that generated when head roll is compensated by a high torsional gain, as observed here (Figures 3 and 4).

More broadly, our data reveals the need to account for torsion to build realistic datasets of visual input during natural behavior. Advances in wearable sensors and computer vision have led to the emergence of large-scale naturalistic datasets combining synchronized gaze, head, and body movements (e.g., Kothari et al., 2020; Ma et al., 2024; Greene et al., 2024). These datasets constitute a powerful resource for the study of natural vision by enabling the reconstruction of visual input during everyday behavior. Here, we show that retinal flow extracted from head and gaze data should be interpreted with caution. Despite our work focusing on the use of retinal flow in locomotion, the influence of torsion in retinal flow shown here has implications for any visual process that computes retinal flow during self-motion, like balance (O’Connor & Kuo, 2009) or object-motion processing (Scherff & Lappe, 2025). Finally, although small in amplitude, the presence of torsion affects eye orientation and, therefore, where on the retina each stimulus falls. This is relevant to studies of visual processes that compute the retinotopic localization of stimuli during complex behavior.

Combined, our work emphasizes the relevance of accounting for the torsional component of eye movements in natural behavior and calls for the refinement of methods capable of delivering accurate and stable measurements of the full 3D eye kinematics in freely moving subjects. Building egocentric and realistic models of visual input is crucial for advancing our understanding of vision (Bonnen, 2023) and will require a fine-grained comprehension of the behaviors that generate them.

## ACKNOWLEDGMENTS

This work was supported by grants PID2023-150081NB-I00 to JLM and PID2023-150883NB-I00 funded by MICIU/AEI/10.13039/501100011033 to CM. AM was supported by grant PRE2021-097688 funded by MICIU/AEI/10.13039/501100011033 and the FSE+ and associated to the Maria de Maeztu project of the Institute of Neurosciences of the University of Barcelona (MDM-2017-0729-21-1). We thank Pablos Marcos-Manchón, Kathryn Bonnen and Gabriel Díaz for helpful discussions. We thank Albert Van den Berg and Eli Brenner for helpful comments on an earlier version of this manuscript. The authors would like to thank Manel Moreno for his help in designing the equipment that made the study possible.

## AUTHOR CONTRIBUTIONS

A.H.M., C.M. and J.L.M. conceived of the experiment; A.H.M. conducted the experiments. A.H.M carried out the data analysis and wrote the manuscript with input from C.M, J.L.M. and J.O.M. All authors reviewed the final version.

## DATA AVAILABILITY

Iris images collected during experiments constitute sensitive biometric data and will not be made publicly available. All other data supporting the findings of this study will be made available upon acceptance of the manuscript.

## METHODS

### Participants

One author (AHM) and nine naive human subjects participated in the study (2 females, 8 males; age: 30.2 ± 5.2 years, height: 1.76 ± 0.10). All gave written informed consent before data collection. Experimental procedures were approved by the University of Barcelona ethics committee (IRB00003099) and are in line with the Declaration of Helsinki.

### Equipment

Subject’s gaze data and head orientation were tracked using a Pupil Labs head-mounted eye tracker (Pupil Labs Neon, Berlin, Germany). Each eye was recorded at 220 Hz by an infrared camera. Head orientation was recorded at 100 Hz by an inertial measurement unit (IMU) also centered - and part of - the eye-tracker (ICM-20948 TDK InvenSense, see Figure 1A top). The IMU was calibrated at the beginning of each recording. Because torsion coding relied on good quality images, we adapted a 30 Hz small camera (Logitech C270, 1280 x 720 resolution) to be mounted on the eye tracker frames together with a dimmable light. A lightweight-custom 3D-printed hinged mounting platform was developed that allowed controlled adjustability while ensuring stability during locomotion (see Figure S1A top). The platform admitted camera positions within −0.5 cm to 1.5 cm (on x axis), −0.5 cm to −2 cm (on y axis) and 3.5 to 5.5 cm (on z axis) with respect to the estimated eye center. The camera focus, the light intensity and the camera and light positions were adjusted according to head, eye and eyelid shape, eyeball light reflection and subject’s light tolerance. The RGB camera was positioned slightly below the eye to minimize occlusions. Finally, the RGB camera frames were manually synchronized with the eye tracker’s clock using a 60 Hz synchronization high contrast video at the beginning and at the end of each trial. The video was presented closely in front of the world camera and was captured by the RGB camera by its reflection in the participant’s eyeball.

### Experimental procedure

Pariticipants were asked to wear the head-mounted device while walking along an outdoor hallway at two different paces and fixating a distant target. At the beginning the distance to the target was 14 m, and participants walked until the target was 4 m away. They did so in a slow and a fast condition. In the slow condition, and were instructed to walk gently as if approaching a target they were curious about. In the fast condition, participants were asked to walk with determination towards the distant target. Average walking speeds across subjects were .58 m/s and .99 m/s for the slow and fast conditions, respectively. Different paces were instructed with the purpose of generating intra-subject walking variability. We avoided explicit instructions for walking fast or slow so that the walking pattern and speed was natural. As torsion detection required good iris images, and natural light is easily reflected by the cornea, all experiments took place at the Faculty of Psychology around 6 pm in an outdoor open hallway with a ceiling (see Figure 1A bottom and Supplementary Videos 1 and 2).

### Signal pre-processing

Head and eye data from the eye tracker were transformed to match X left, Y up and Z forward axis. Eye data was further rotated 12 degrees to correct for the mismatch between IMU and EYE frames in the Neon eye tracker. Head orientation and velocity data was obtained from the IMU at subject’s forehead. In a few trials, unexpected saccades were made to large, eccentric positions and were eliminated from analysis. Eye movement data from the eye tracker was smoothed using a 4th-order zero-phase Butterworth low-pass filter with a cutoff frequency of 40 Hz to remove high-frequency sensor noise. All data was down sampled and matched to our RGB camera where iris features were tracked. Missing feature rotation data created by blinks or blur were interpolated using Akima piecewise cubic interpolation (blinks: 3.1%, blurred eye feature: 0.98%) and the resulting signal was smoothed with a 4th-order zero-phase Butterworth low-pass filter at 10 Hz.

### 3D and 2D frames of reference

Four 3D frames of reference will be relevant to our 3D data analyses: world, locomotion, head and eye (see Figure 1A top). The world frame of reference is used to report head orientations by the head-mounted eye tracker (with the Z axis pointing to the magnetic North Pole). The locomotion frame of reference is aligned to the subject’s trajectory and fixed in the world. As participants walked in a straight line, we computed this frame of reference by interpolating the head orientation quaternion data. The head and eye frames of reference moved with the partiicipant. Again, as participants walked in a straight line, stabilizing their gaze on a target in front of them, the head’s and, more specifically, the eye’s z axis was almost parallel to the locomotion z axis.

For the gaze-related analyses, three 2D frames of reference will be relevant: camera image, stabilized image and world (see Figure S2B). Gaze camera image positions are raw pixel data that describe gaze on the image and are independent of how subjects are oriented in space: changes in gaze on this frame are changes of eye orientation in the head. The stabilized image frame reports gaze positions with respect to an external reference. To do this, we used automatic computer vision algorithms (Karaev et al, 2024) to track salient features near the target throughout the video (see Supplementary videos 1 and 2). Frame-by-frame post-processing corrected for features that were offset on a few frames. Gaze and wall-feature positions were first undistorted and registered, frame-by-frame, to a common reference frame via a rigid-body (rotation-and-translation-only) transform, fitted to the tracked wall features. In both these frames, pixel values correspond almost linearly to visual angles within close proximity to a central target, consistent with gaze behavior in our data. To illustrate the distinction between these frames: if a subject rotates the head to the left while fixating the target, the eyes rotate to the right, shifting gaze position in camera image coordinates - but gaze position in the stabilized frame remains largely unchanged. Finally, we obtain world 2D frames by rotating, translating and scaling camera images to the same common reference frame and report gaze in an external fixed frame where pixels are related to world distances.

### Torsion estimation

In the context of natural behavior pure rotations around the line of sight, or any other direction, are infrequent (see Figure 1B right). Here we are interested in the eye-centered (intrinsic) rotational displacement components along the line of sight and we will report torsion as a velocity. Velocity is independent of sampling rate, making velocity-based measures directly comparable across recording systems (as in Tweed & Vilis, 1990). In other words, given a small rotation, described by the rotation vector dr = [dr_x_, dr_y_, dr_z_], happening between two frames at dt (33 ms in our data), dr_z_/dt represents our measure of torsion (being the eye Z axis, its line of sight). As it refers to eye intrinsic axes, this measure is not affected by the constant changes in line of sight during locomotion and is directly related to the rotational component of retinal flow.

In three dimensions, the rotation of features around the line of sight is directly related to the torsional component of the eye’s rotation velocity vector. Camera projections, however, create artifacts that depend on camera position and the eye axis of rotation. We exploited two properties that make estimating torsion from a 2D signal tractable. The first is that gaze direction can be estimated at every interval. Knowing the eye’s gaze direction at each frame and between two consecutive frames does not, by itself, specify the eye’s three-dimensional rotation between those frames - but it constrains the problem substantially. Second, competing and physiologically motivated eye rotation scenarios can be simulated to produce distinguishable, non-overlapping predictions for iris feature rotations measured on the camera image. No torsion and Listing’s torsion can be computed from two consecutive gaze positions and VOR-like rotations can be simulated from gaze position and head velocity data. Notably, if gaze stabilization is precise, all scenarios vary only on their torsional component. Perfect gaze stabilization implies the eye stabilizes the head in the horizontal and vertical axis, leaving only torsion as the unknown variable. In this case, simulations then consist of 100 VOR scenarios varying just on the torsional component (gains 0 to 1), being two of those scenarios, the torsion-free (gain = 0) and the Listing scenario. In the alternative gaze vector calculations, 3D gaze is extracted from azimuth and elevation and will not always reflect perfect head compensation on the horizontal and vertical axis. In this case, 99 scenarios represent the eye compensating for the head data with different torsional gains (.01 to 1). Torsion-free and Listing scenarios are calculated from the gaze data and do not overlap with any simulated VOR gain scenario. In these calculations we then have 101 scenarios in total. As mentioned above, the alternative ways of estimating 3D gaze vectors did not affect the conclusions of this work.

We designed a lightweight device that mounts on the head-mounted Neon eye tracker frame (Pupilabs, see Fig S1A) and applied automatic tools to track multiple iris features that were visible throughout the whole trial (Karaev et al, 2024, see supplementary videos 1 and 2). 5 to 10 features distributed around the pupil were selected for each subject. Pupil location was detected using open-source python libraries from Pupil-Labs: https://github.com/pupil-labs/pupil-detectors. A frame-by-frame post-processing was done to discard noisy features and manually correct stable features that were off in just a few frames (same with pupil). The coder did not have access to subject’s data that could bias the coding process (i.e., head roll). Feature rotation was computed as the mean change in angular position across iris feature relative to the pupil center (empirical feature rotation).

Torsion was estimated from a simulation-based framework linking eye kinematics to their camera projection, in several steps. First, for each frame, eye gaze was used to render the pupil and six iris features – radially distributed around the pupil. Listing’s plane was assumed to be orthogonal to the line of sight in the forward-facing position. Then, the rotational displacement between two frames was calculated to render the new position of the iris features. For the torsion-free scenario, the rotational displacement was computed from the cross and dot product of their gaze vectors. For the Listing’s scenario we first calculated, for each frame, the orientation defined by Listing’s law given each gaze position. We then calculated the extrinsic fixed-axis displacement in head coordinates that takes one frame to the next by:

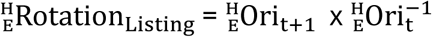

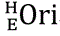 stands for the orientation matrix of the eye in the head on a given frame. For the VOR scenario, the rotation between frames was extracted from the head data. One hundred VOR scenarios were simulated, each with a value from .01 to 1 on its torsional gain. As mentioned above, for all scenarios, the compensatory eye rotational displacements were calculated and used to rotate the iris features between pair of frames.

Second, using a pinhole camera model, we projected the 3D feature points (P) onto the image plane via the estimated camera intrinsic parameters:

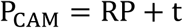

**R** and **t** are the camera rotation and translation and P_CAM_ are the resulting coordinates in the camera plane. The effective focal length f was derived from the physical camera sensor width w and the corresponding field of view (FOV) angle **α**. For every pair of consecutive frames – each scenario and each camera - the mean change in angular position across iris feature relative to the pupil center was computed, as we did with our empirical data. Variables and scripts to run the simulations for all subjects and trials are available (see Data availability).

As a final step, to quantify the agreement between the reference and empirical signals, we used the concordance correlation coefficient (CCC) proposed by Lin (1989):

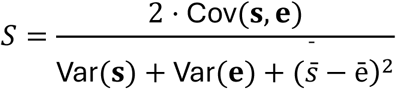

where Cov(**e**, **s**) is the covariance between the simulated **s** and the empirical signal **e**, Var denotes variance, *s̄* and *ē* are their respective means. The denominator decomposes mismatch into three terms: variance of the simulated signal, variance of the empirical signal, and squared mean bias. The score ranges from −1 to 1, where *S* = 1 indicates perfect agreement. By computing S for all pairs of signals, we found the VOR gain value that best matched our empirical data. Finally, the temporal relation between the empirical feature rotation data and the best-fitted-VOR scenario was quantified using the normalized cross-correlation function (Pearson) at lags up to ±400 ms.

### Retinal flow and curl estimation

Retinal flow structure is a direct consequence of the translation and rotation of the eye with respect to the environment and the geometry of the surrounding space. Assuming a static environment, the flow can be decomposed into its translational and rotational components (Longuet-Higgins & Prazdy 1980). In our data, because subjects stabilized gaze at a distant target, the structure of flow is greatly simplified. As subjects walked while stabilizing gaze on a distant target, translational flow would resemble expansive patterns and non-compensated head roll components would result in flow curl (as simulated by Tweed et al, 1994b). Linear translation velocity was available from the trial length and duration (slow trial mean: .58 m/s; fast trial mean: .99 m/s). However, because we were specifically interested in the rotational component of flow generated by head rotations, we used a fixed linear speed of 1 m/s to allow direct comparison across subjects and translations of the head in the lateral and vertical directions were not modelled.

Retinal flow was calculated using the equations relating 3D motion to optical flow (Longuet-Higgins & Prazdy 1980; Heeger & Jepson 1992) on the no torsion and VOR scenario (its best fit gain) by modelling eye kinematics from both scenarios on a simulated infinite floor. The curl of the flow field **F** = (*F_x_*, *F_y_*) was then computed at each grid point as:

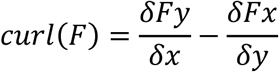

Curl magnitude was defined as the peak absolute curl value across the visual field, providing a single scalar summary of the maximum rotational flow intensity per frame.

