## Supplementary information for "Torsion in motion: the visual system as a three-axis gimbal"

### Three-dimensional eye rotations, torsion and Listing's law

#### *Eye and head rotational degrees of freedom*

Fully specifying the orientation or rotational displacement of a rigid body in space – like the head or eye - requires, each, three degrees of freedom. Regardless of the choice of parametrization, the coordinate frame or the kinematic aspect being studied, these three degrees of freedom always capture some horizontal, some vertical, and some forward component of rotation, the latter often referring to a rotation about the forward or visual axis. Head degrees of freedom are often referred to as yaw, pitch and roll. These terms can be interpreted in two ways. Thay can refer to the choice of representation of head orientation as the sequence of ordered rotations (Euler angles), or – as here - as mere labels to an axis of reference: X or pitch, Y or yaw and Z or roll. We note that these risks conflating these two distinct uses of these terms However, we retain this terminology for consistency with literature and for readability but flag the distinction explicitly to avoid ambiguity.

Both for the eye and the head horizontal and vertical axis have received most attention (Pozzo et al., 1990; Moore et al., 1999; Moore et al, 2001) as horizontal and vertical rotations, but not roll or torsion, shift the field of view, a central behavior for visual processing and cognition. At small steps, the component of any rotation along the eye's line of sight does not alter the displacement at the center of the foveal image. This is illustrated in Figure 1B for the eye. The rotational velocities ($\omega$) differ in their torsional component, with $\omega_2$ being orthogonal to the line of sight and therefore having no component about it. The velocities in the fovea (v), however, are the same. This implies that only two components can be enough to describe gaze stabilization in head free-conditions.

How the eye rotates in the head has been largely studied in controlled laboratory conditions. Donders' and Listing's law represent an elegant example of how sensorimotor processes can constrain degrees of freedom, both in orientation and displacement. Some confusion, however, has emerged about what Listing law implies for eye rotation. These laws describe orientation during fixations, saccades and pursuit and have consequences for eye rotational displacements. However, as we will see below, in most cases, unless the eye is close to its primary position or making saccades or pursuit from or towards its primary position, even during saccades all three degrees of freedom are needed to fully represent eye orientations or rotational displacements.

#### *Donder's and Listing's law: constraints in eye orientation*

Two foundational laws describe how the eyes are oriented in space when directing gaze to a target and set the stage for the study of eye movements. Donder's law states that for any gaze direction, the eye always assumes the same 3D orientation, no matter where the eye came from. Listing specifies that there is one position (primary) from which you can uniquely describe all others by a rotation that lives in a plane (Listing's plane) that is normal to the primary position (Ferman et al,

1987b). Two relevant implications follow. If all orientations can be described from primary position by rotations around axes in one plane, all eye orientations can be described with two parameters. Secondly, given that the plane is normal to gaze direction on primary position, all other eye orientations from there can be described by rotations with no component along the line of sight or no torsion (van den Berg, 1995).

The description of eye orientation from a reference position and how it moved through space to get there are fundamentally distinct problems (Haslwanter, 1995). The empirical study of saccades and smooth pursuit reveals, however, another elegant geometrical constraint. No matter where the eye rotates from, all the possible rotations from its initial orientation can be described by axes on one plane (Tweed et al 1990, called displacement planes, see Figure S1C). The orientation of each displacement plane in any position is determined by the gaze vector in that that position and by the primary position. More precisely, their orientation is normal to the vector that bisects primary position and the current gaze direction (reason why they are also called half-Listing planes, see Figure S1C). As an example, if the eye is looking 30 degrees to the right, the displacement plane will be normal to the bisector vector 15 degrees to the right. When the eye is in primary position, the bisector corresponds to the gaze vector in primary position itself (angle=0). The primary position is then the unique position in which that plane is normal to the gaze direction. That is, Listing's plane is the unique case where the displacement plane is normal to the gaze position, and therefore, to reach any orientation from the primary position the eye rotates with no torsion. Two new - and apparently incompatible - implications arise. They apply to saccades or pursuits occurring between secondary or tertiary positions, with the specific exception of the second implication not applying to rotations towards or away the primary position. First, at any step throughout its journey, the eye is oriented in a way that can be described by a rotation whose axis lives in Listing's plane (perpendicular to primary and fixed in the head). Second, at any step the eye is rotating around an axis that has components along the line of sight (Tweed & Vilis, 1990). The larger the distance from primary position, the larger the tilt of the half-listing or displacement plane, the larger its component around the line of sight. In other words, to follow Listing's law, the eyes must rotate with some torsion, that grows with eye eccentricity and is defined by the tilt of its displacement plane.

*Torsion: a conceptual challenge*

A rotation of the eye about the line of sight when gaze is directed forward provides an indisputable example of ocular torsion. What is torsion outside this specific context is much less evident. The different choices of parametrization - rotation matrices, quaternions, rotation vectors, or Euler angles -, different frames of reference - world, head and eye -, and the aspect of eye kinematics under consideration - orientation or velocity - have led to divergent definitions of torsion. The word torsion has been used to describe the rotation around the line of sight as a response to pure head roll (Collewijn et al, 1985), the angular velocity component along the line of sight (Tweed & Vilis,1990) and the tilt of the vertical axis of the eye with respect to gravity (Khazali et al., 2020). To make it even more challenging, the multiple ways in which 3D kinematics can be represented (Euler angles, rotation matrixes, rotation vectors, quaternions; Haslwanter, 1995) render different torsional components that are a product of the choice of representation (false torsion, Carpenter, 1988). What is torsion, therefore, needs to be clearly defined to avoid confusion.

*Torsion in natural behavior: a technical challenge*

The non-invasive measure of the eye's full 3D kinematics in natural behavior remains elusive. The eye pupil properties (geometry, position, and high contrast) that enable accurate tracking of horizontal and vertical eye movements are not informative about rotations about the line of sight. Extracting torsion from images has mainly focused on iris detection which requires dealing with artifacts generated by eyelid occlusions and distortions of the iris generated by eccentric positions of the eye (see Otero-Millán et al, 2015 for a review). In the lab, advances in iris pattern recognition algorithms show promising results within non-eccentric positions of the eye (+/- 10 degrees vertically and +/- 20 degrees horizontally, see Otero-Millán et al, 2015). Measuring torsion outside the lab adds the challenge of designing an eye tracker with little occlusion of the field of view, short focus distance, light weight, that deals with shadows and reflections on the cornea generated by non-controlled lightning conditions that change along the environment and with gaze shifts.

**Eye stabilization in natural behavior**

Our setup was specifically designed to show the pervasiveness and functional relevance of torsion outside the lab. In the study, subjects fixated a stationary target at a minimum distance of four meters, a frequent behavior in natural locomotion (Holland et al, 2002; Hart & Einhäuser 2012). At this distance and position, several factors that complicate the interpretation of eye movements in natural behavior are minimized. First, a technical facilitation – good resolution for iris tracking can be largely affected by shadows as eyes move. The small range of eye movements while stabilizing at the front enables the delimitation of an area where this is possible. Shadows and reflections on the cornea would make wider movements harder to track by our approach. Second, eye rotations near primary position (median head displacements around 2 degrees) simplifies the geometric interpretation of torsion. Camera's viewing axis remains approximately aligned with the eye's optical axis. Third, vergence is minimal — with standard interocular separation each eye deviates less than one degree from parallel at the end of the trial (0°43'). Fourth, translational head movements during walking, both vertical bob and lateral sway, create angular retinal demands that increase with target proximity and can be compensated by eye or head rotations; Moore et al. (1999)

showed that for distant targets (more than two meters) the angular VOR carries the stabilization load. At these distances, head rotations and translations acted as perturbations to gaze rather than contributions to it (Moore et al, 1999; Moore et al, 2001) - a distinction that would not hold were subjects fixating a target on the floor ahead. In this case, pitch head rotation would move gaze toward rather than away from the target. Finally, the deviation between required eye counter rotation and actual head rotation angle, which arises from the offset between the eyes and the head's center of rotation, becomes negligible with target distance.

What reaches the retina during natural behavior is the consequence of a sophisticated sensorimotor process involving the whole body, mediated by behavioral goals and extending well beyond the factors considered here (Ibbotson et al., 2005; Lambert et al., 2023; Moore et al., 2001; Wibble & Pansell, 2024). Even within fixations eyes are known to move in head-free conditions (Poletti et al., 2015), setting a lower bound on what perfect stabilization means in practice. A fuller account of eye stabilization during locomotion will require simultaneous measurement of all six degrees of freedom of head motion over time, a 3D model of the surrounding environment and a clearer understanding of the role of fixations in natural behavior (Lappi, 2016).

**3D gaze vectors and gaze stabilization**

Gaze parameters extracted from the head-mounted eye tracker (azimuth and elevation) describe gaze position in the image. The gaze vector in the head was estimated in two alternative ways. On one hand, gaze was computed from head orientation data assuming perfect gaze stabilization - an assumption supported by prior reports (Grossman et al. 1989). On the other hand, gaze was computed from azimuth and elevation on the camera image. At the viewing distance used in this study (14 to 4 meters), azimuth and elevation provided by the head-mounted eye tracker with respect to the scene camera diverges from the eye gaze vector in the head by negligible amounts. Given a 30 mm eye-to-camera offset, this divergence is largest at the closest approach distance (4 m), where the scene camera's forward direction (0°) differs from the true eye-to-target vector by approximately 0.43°. The gaze vectors obtained were then rotated to correct for any offset between the eye tracker origin for azimuth and elevation, and the baseline gaze position throughout the trial, calculated as the geometric median of all gaze vectors. For both methods of estimating gaze, the 5-second window with best stabilization was selected for each trial.

Gaze stabilization was computed from gaze data on the stabilized 2D frame discussed in the 3D and 2D frames of reference on the Methods section. Within each window, gaze accuracy was defined as the mean angular distance between gaze and the target; gaze precision was defined as the root mean square (RMS) angular distance between gaze and its own median within that window, capturing internal gaze consistency independent of target location. Precision was used as our measure of gaze stability to select the window for analyses for each trial. All results shown in this study were consistent between the two gaze-in-world estimation methods across these windows. As a further check, we ran all analyses on the 5-second window within each trial where the two gaze estimates described above – from azimuth/elevation and assuming perfect stabilization - were most correlated with one another, rather than selecting based on gaze stability on the world image. Results converged on the same conclusions. In the main text, we report results using gaze extracted under the perfect-stabilization assumption on the 5 s window with best stability extracted from gaze in the image (Figure S3 shows results for the alternative estimations).

**Camera uncertainty**

To assess whether uncertainty in the estimated camera position and orientation could account for our results, we generated 560 plausible camera configurations per subject, sampled around each subject's estimated camera (translational offsets up to 1 cm in every direction and roll offsets up to 10°). At the measured camera-to-eye distance (~4.1 cm), a 1 cm positional offset orthogonal to the viewing direction corresponds to an angular deviation of approximately 13.6°. Gain was recomputed for every subject × condition × camera combination (11,220 values total). A linear mixed-effects model (gain ~ condition + (1|subject) + (1|camera)) estimated that camera identity accounted for a negligible share of the total variance in gain (estimate ≈ 0%), compared to ~54% attributable to between-subject differences, indicating that our results are not sensitive to our estimated uncertainty in camera placement (Figure S2D).

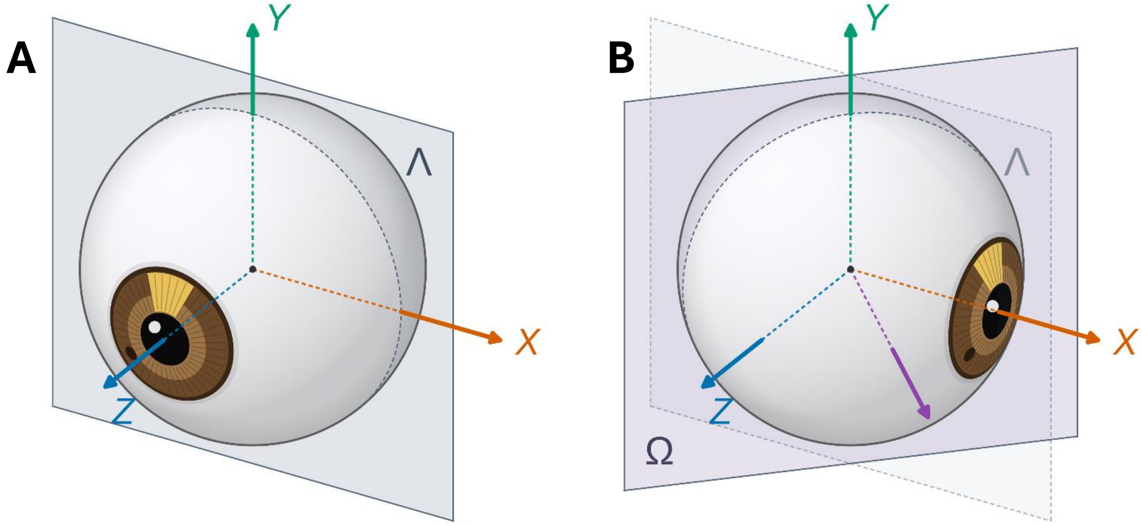


**Fig S1.** *A*. The figure shows the eye in primary position. Primary position is fixed in the head and approximately aligned with the forward direction (Z axis). Y denotes the vertical axis. As oculomotor research shows, given any initial position, the eye rotates in a plane that is normal to the vector that bisects the initial gaze vector and the gaze vector in primary position. When the eye is in primary position as in the figure, the two vectors are the same and the eye will rotate around an axis that lies in the plane normal to primary position itself, i.e., in Listing's plane (Λ, also fixed in the head). In other words, from the primary position the eye rotates without components along the line of sight or torsion. *B*. The figure shows an extreme case in which the initial gaze direction is 90° to the left (overlapping the red x axis) - beyond the physiological range but chosen for clarity. The violet vector (45 degrees to the left) bisects the angle formed by the eye's current gaze vector and the gaze vector at primary position. Consequently, the violet vector is the normal vector to the plane (Ω) that contains the candidate rotational axis that can be carried out from this initial position. Ω is then the displacement - or half-Listing's - plane for this gaze position. In other words, to make a saccade from here, the eye will rotate around a vector contained in Ω. Listing´s plane (Λ) is shown for comparison

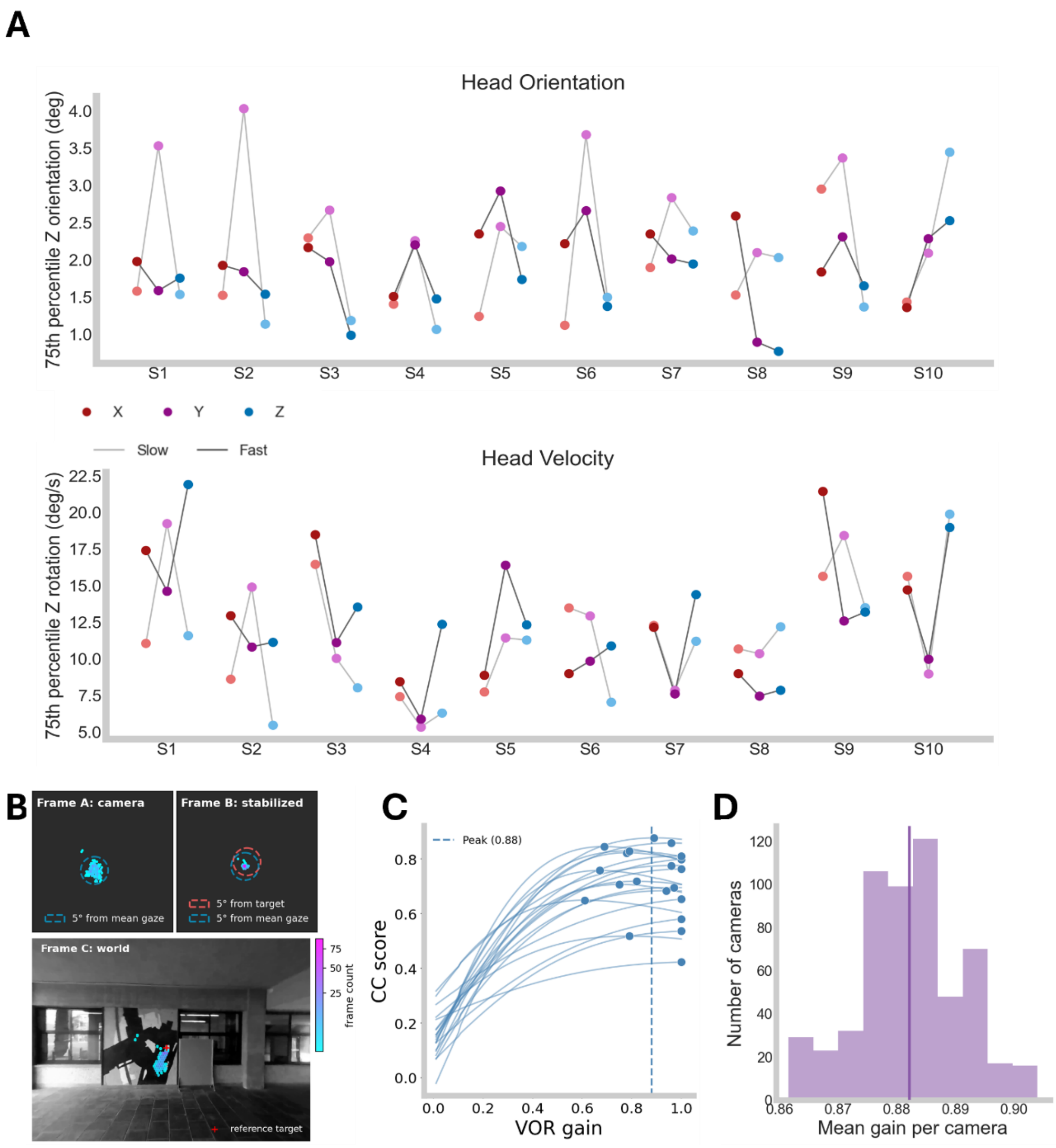


**Figure S2. A.** *Top*. For each subject, the 75th percentile of each component of the head's orientation vector is displayed for both the slow (light colors linked by grey lines) and fast (darker colors linked by black lines) trials. *Bottom*. Same as A but with angular velocity vector components. **B**. Reference frames associated with gaze during natural behavior. Frame A is the position of gaze in the camera image (top left). Frame B is a stabilized version of Frame A, without scaling (see 3D and 2D frames of reference section). Frame C shows gaze in the world. The last frame of the trial was selected as the reference where to map gaze on Frame C. **C.** Gain fit curve for each subject and trial. Each dot represents the VOR gain value that yields the maximum CCC score for one trial, and which defines the torsional component of the eye that best explains the empirical eye data. **D.** Effect of camera position on the fitted gain values. Each value in the histogram represents the mean across subjects and trials for each of the 561 cameras. Vertical line represents the corpus mean.

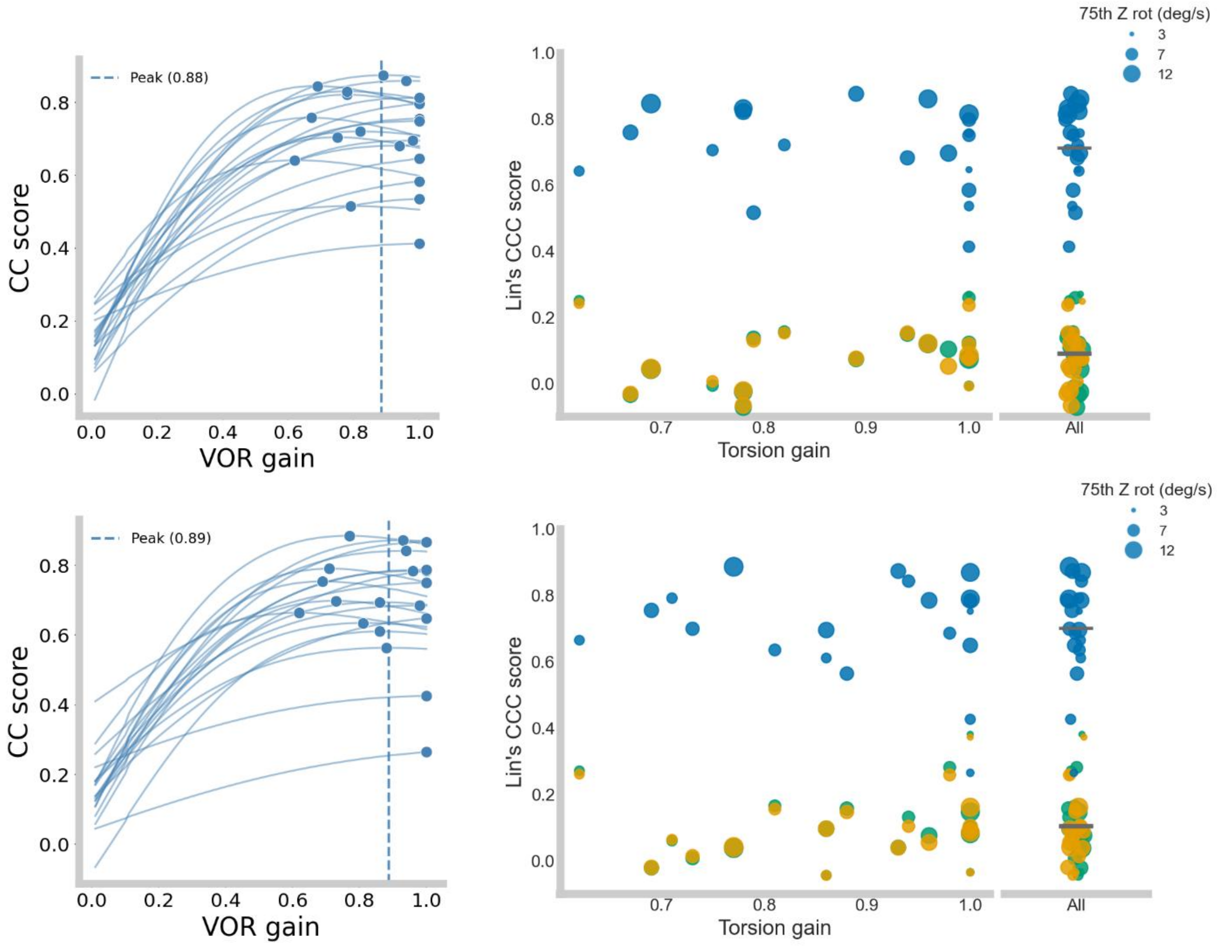


**Fig S3.** *Top.* Both figures show the results obtained for figures S2C and 1C right if 3D gaze vectors in the head were calculated from the azimuth and elevation image gaze data from the head-mounted eye-tracker. *Bottom***.** Both figures show the results obtained for figures S2C and 1C right if 3D gaze vectors in the head were calculated from the azimuth and elevation image gaze data from the head-mounted eye-tracker and gaze stability was not computed from gaze in the image but from the correlation between 3D gaze and head rotations (see section 3D gaze vectors and

**Supplementario Video 1.** Empirical data extracted from one trial. Top signals show head roll velocity (left) and 2D feature rotation velocity (not torsion, right). Video on the lower left shows the egocentric view from subject's forehead. Target (red), gaze (green) and wall features (yellow) are tracked. Video on the lower right is blurred and showed in greyscale and shows the eye with the selected tracked features (blue).

**Supplementario Video 2.** Empirical data extracted from one trial. Top signals show head roll velocity (left) and 2D feature rotation velocity (right). Video on the lower left shows the egocentric view from subject's forehead. Target (red), gaze (green) and wall features (yellow) are tracked. Video on the lower right is blurred and showed in greyscale and shows the eye with the selected tracked features (blue).
